# Cell cycle-dependent recruitment of FtsN to the divisome in *Escherichia coli*

**DOI:** 10.1101/2021.12.09.472041

**Authors:** Jaana Männik, Sebastien Pichoff, Joe Lutkenhaus, Jaan Männik

**Author notes:** Address correspondence to Jaan Männik;, Joe Lutkenhaus.

## Abstract

Cell division in *Escherichia coli* starts with the formation of an FtsZ protofilament network in the middle of the cell, the Z ring. However, only after a considerable lag period do the cells start to form a midcell constriction. The basis of this cell cycle checkpoint is yet unclear. The onset of constriction is dependent upon the arrival of so-called late divisome proteins, among which, FtsN is the last arriving essential one. The timing and dependency of FtsN arrival to the divisome, along with genetic evidence, suggests it triggers cell division. In this study, we used high throughput fluorescence microscopy to quantitatively determine the arrival of FtsN and the early divisome protein ZapA to midcell at a single-cell level during the cell cycle. Our data show that recruitment of FtsN coincides with the initiation of constriction within experimental uncertainties and that the relative fraction of ZapA/FtsZ reaches its highest value at this event. We also find that FtsN is recruited to midcell in two distinct temporal stages with septal peptidoglycan synthesis starting in the first stage and accelerating in the second stage, during which the amount of ZapA/FtsZ in the midcell decreases. In the presence of FtsA*, recruitment of FtsN becomes concurrent with the formation of the Z-ring, but constriction is still delayed indicating FtsN recruitment is not rate limiting, at least under these conditions. Finally, our data support the recently proposed idea that ZapA/FtsZ and FtsN are part of physically separate complexes in midcell throughout the whole septation process.

**Importance:** In *E. coli*, FtsN has been considered a trigger for septal wall synthesis and the onset of constriction. While FtsN is critical for cell division, its recruitment kinetics to midcell has not been characterized. Using quantitative high throughput microscopy, we find that FtsN is recruited to midcell in two temporal stages. The septal cell wall synthesis starts at the first stage and accelerates in the second stage. In the presence of an FtsA mutant defective in self-interaction, recruitment of FtsN to midcell is enhanced, but constriction is still delayed. Our results shed new light on an essential but not rate-limiting role of FtsN in *E. coli* cell division and also support the view that ZapA/FtsZ and FtsN are part of physically separate complexes in midcell throughout the division process.

## Introduction

Cell division in bacteria requires the regulated assembly of several dozen divisome proteins that collectively initiate and guide the formation of the constriction/septum, which divides the mother cell into two daughters (1, 2). In *Escherichia coli*, the assembly of the essential divisome proteins involves at least two stages (3, 4). During the first stage, the tubulin-like GTPase FtsZ polymerizes into dynamic protofilament assemblies (5) that are attached to the inner leaflet of the cytoplasmic membrane by the membrane tethers FtsA and ZipA (6). These assemblies are also decorated by the non-essential Z-ring associated protein, ZapA (3, 5, 7), and potentially with several less conserved and less abundant Zap proteins (ZapC, ZapD) (8–10). This initial complex forms a highly dynamic discontinuous ring-like structure, referred to as the Z-ring, composed of FtsZ protofilaments which treadmill around the division plane (11–13). FtsEX also arrives at this stage and is required for recruitment of proteins during the second stage (14, 15).

During the second stage, the remaining essential divisome proteins are recruited to midcell to complete the assembly of the mature divisome (1, 2), which then starts to synthesize the septal cell wall. Dependency studies indicate a sequential recruitment of these divisome proteins with the order FtsK<FtsQ<FtsB-FtsL<FtsW-FtsI<FtsN (1, 2). FtsN is the last of the essential proteins to arrive, and it has been postulated to be the trigger protein that leads to the onset of constriction (16–21). FtsN is an integral membrane protein (22), with a short cytoplasmic domain (FtsN^cyto^), a single transmembrane helix followed by a large periplasmic domain (23). The periplasmic domain is further divided into two subdomains: an essential domain (FtsN^E^), which has been implicated in triggering septal peptidoglycan (PG) synthesis (17, 24), and a C-terminal SPOR (septal peptidoglycan binding) domain, which interacts with PG strands denuded of peptide linkages by amidases (16, 25, 26).

The recruitment of FtsN to midcell is complex, requiring FtsA, FtsQ, FtsI and likely all upstream proteins (1, 27, 28). It has been reported that a small amount of FtsN is recruited to the division site before the start of cell wall synthesis through a cytosolic interaction between FtsN^cyto^ and FtsA at the Z-ring (29–32). This initial recruitment of FtsN, along with ZipA, is proposed to interact with the PG synthases PBP1a and PBP1b to initiate a pre-septal phase of PG synthesis (33). Once the constriction starts, more FtsN is recruited via its SPOR domain that binds to denuded glycan strands produced by activated amidases (16, 34). Since the denuded strands are formed following the onset of septal cell wall synthesis, the recruitment process of FtsN via its SPOR domain is thought to be self-enhancing (also referred to as the septal PG loop) once constriction starts (16, 22, 25).

Current models on how FtsN initiates the onset of constriction envision a two-pronged mechanism (17, 35–37). In this model FtsN^E^ causes a conformational change in FtsQLB in the periplasm that activates FtsWI (36). A parallel activation pathway in the cytoplasm involves FtsN^cyto^ activating FtsA (30–32). The FtsA mutant R286W, with reduced self-interaction, is thought to mimic an active state of FtsA and will be denoted here also as FtsA*. FtsA* acts directly on FtsW in the cytoplasm (38). Under physiological conditions these pathways synergize to activate FtsWI. However, mutations that hyperactivate either of the two pathways are capable of leading to cell constriction in the absence of the other (1, 17, 35).

While FtsN is essential for cell division and potentially a trigger for it, the kinetics of its recruitment to the divisome has not yet been determined. In particular, the question of whether FtsN recruitment is rate-limiting for the onset of constriction has not been addressed. All existing data on FtsN recruitment to the divisome originates from static cell studies or imaging studies where short time-lapse series have been taken. Here we study FtsN dynamics at the individual cell level throughout the cell cycle and determine its recruitment kinetics to the divisome. Our high-throughput studies show that while the recruitment of ZapA/FtsZ to midcell is gradual, recruitment of FtsN is abrupt and occurs on average about a quarter of the cell cycle after a persistent Z-ring forms. Within minutes of FtsN recruitment, if not faster, the onset of constriction starts indicating that the septal cell wall synthesis is tightly linked to the arrival of FtsN to midcell. In the presence of FtsA*, FtsN arrives at midcell as the Z ring forms but constriction is not immediately initiated. Under these conditions FtsN arrival is not the rate-limiting component of the divisome that determines the timing of the onset of constriction. We also find that FtsZ protofilament condensation is not required for FtsN recruitment and the onset of constriction as has been proposed for *B. subtilis* (39, 40). However, a fraction of ZapA/FtsZ at the midcell in the ring plateaus at its peak value. Furthermore, our data show that the recruitment of FtsN to midcell occurs in two distinct stages. During both stages, cells constrict, but the speed of septal closure increases in the second stage, which starts when ZapA/FtsZ numbers at midcell start to decrease. Our data also lends strong support to the idea that FtsN and ZapA/FtsZ are part of spatially separate complexes throughout the division process.

## Results

### FtsN accumulates rapidly at midcell about a quarter of the cell cycle time after the formation of a persistent midcell ZapA/Z-ring

To investigate the recruitment kinetics of FtsN to the divisome in live cells, we used quantitative fluorescence microscopy of a functional N-terminal fusion of Ypet to FtsN (Ypet-FtsN) (41). We studied the kinetics of FtsN relative to the formation of the Z-ring. As a proxy for the Z ring, we used a functional C-terminal fusion of mCherry to ZapA (ZapA-mCherry) (34). ZapA is a highly conserved early cell division protein that binds to FtsZ, although not essential (42). We found that the ZapA fraction at midcell is precisely the same as the FtsZ fraction at midcell throughout the cell cycle (SI Fig. S1). The same partitioning ratio of FtsZ and ZapA at midcell indicates that they bind to each other at a fixed stoichiometric ratio irrespective of whether FtsZ protofilaments form transient assemblies or are part of a mature divisome that synthesizes septal PG. Altogether, these data show that ZapA-mCherry acts as a faithful reporter for FtsZ. We imaged cells in mother-machine microfluidic devices under moderately fast and slow growth conditions. Cells grew in these devices in steady-state conditions with doubling times of *Td* = 78 ± 28 *min*(*mean* ± *SD*) and *Td* = 143 ± 45 *min*, respectively. These doubling times and cell lengths were comparable to the parent strain without the Ypet-FtsN and ZapA-mCherry labels indicating the fluorescent tags did not affect cell parameters (SI Table S1). Note that all measurements were carried out at 28°C where the growth rate is expected to be about two times slower than at 37°C (43). At the single-cell level, the mid-cell accumulation of ZapA at the slow growth rate increased gradually as a function of cell cycle time (Fig. 1A,B, top), consistent with the previous reports which monitored FtsZ (5, 44). The overall increase in the ZapA amount at midcell was interspersed by large fluctuations in ZapA numbers in the first half of the cell cycle, reflecting the appearance and disappearance of FtsZ transient assemblies (5). FtsN did not appear to participate in these transient assemblies, and its midcell accumulation occurred at later times of the cell cycle and showed a much more rapid increase (Fig. 1A,B, middle). The initial increase in the amount of FtsN at midcell was accompanied by an increase in the mid-cell phase signal (Fig. 1A,B, bottom), indicating that the onset of constriction started at about the time FtsN midcell recruitment as concluded before (16, 18, 20, 41). Thus, FtsN showed an abrupt accumulation at midcell compared to FtsZ and ZapA. Qualitatively similar behavior was also observed in moderately fast-growing cells (SI Fig. S2).

**Fig.1.**
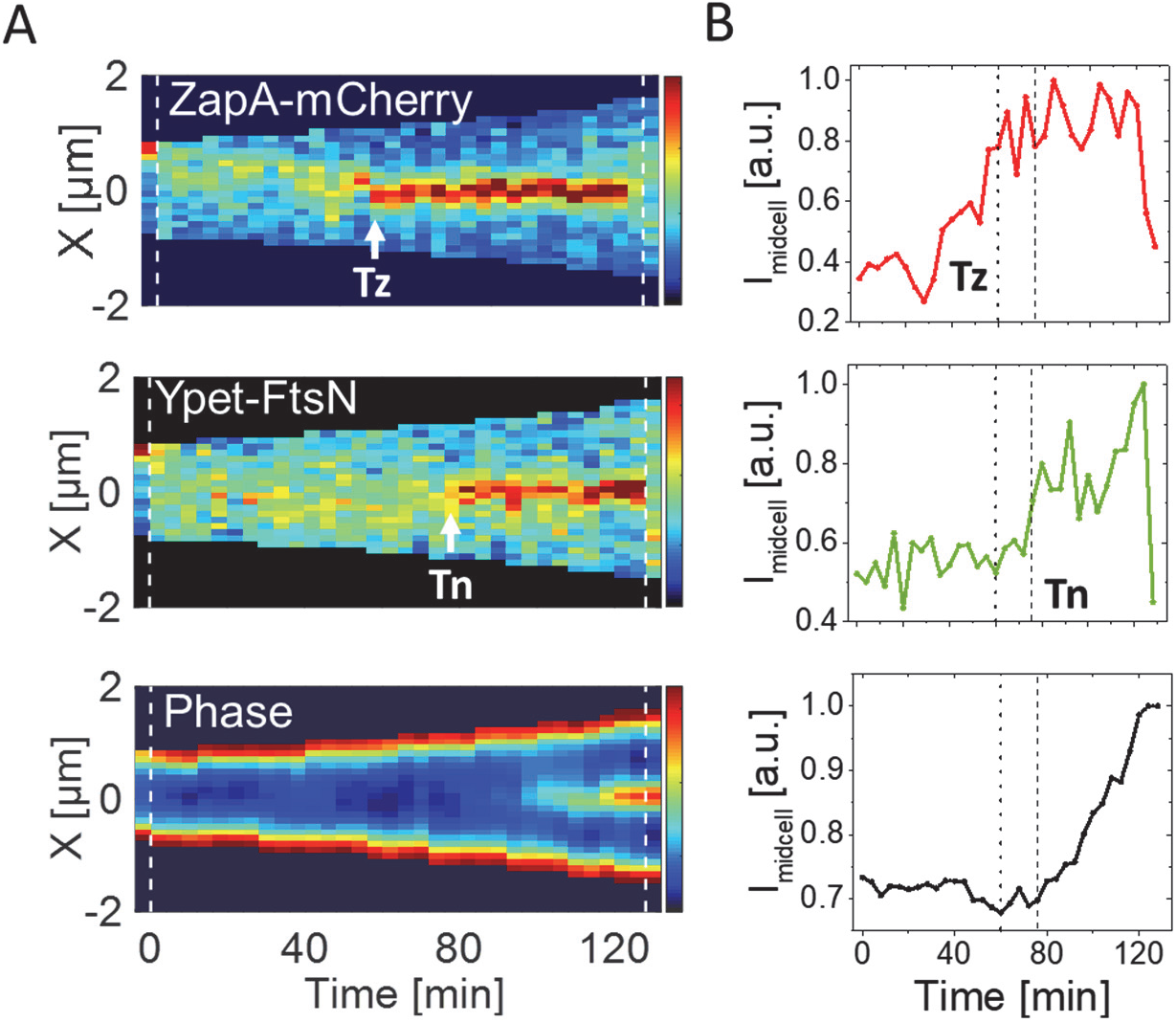
Accumulation of ZapA and FtsN at midcell at the single cell level. **(A)** Kymographs of fluorescent and phase signals for a representative cell grown in M9 glycerol-TrE medium. Red corresponds to high and blue to low-intensity values. Black marks regions outside the cell. Dashed vertical lines indicate cell division events. The arrows indicate event timings as determined by an automated algorithm (see Methods). The timing of the persistent Z-ring is denoted by *Tz* and the onset of FtsN recruitment by *Tn*. **(B)** Midcell intensity traces of ZapA-mCherry (top), Ypet-FtsN (middle), and phase signal (bottom) for the cell shown in the kymograph. The intensity traces were collected from about a 0.5 Ͱm wide band in the cell middle. The dotted line corresponds to *Tz* and the dashed line to *Tn* in all panels. “a.u.” stands for an arbitrary unit. Note that the increase in the phase signal starts at *Tn*.

We then determined the timings of Z-ring formation (*Tz*) and the onset of FtsN accumulation at midcell (*Tn*) from time-lapse images (Fig. 1A) using an automated algorithm (Methods). We refer to time *Tn* as the time when an N-ring is first observed and *Tz* indicates the time when a persistent Z-ring forms, i.e. ZapA/FtsZ protofilament assemblies that are continuously present until shortly before cells separate when the Z-ring disassembles. There are some uncertainties in distinguishing persistent FtsZ assemblies from transient ones (5) in these time-lapse measurements. However, a fully automated and traceable approach allowed us to determine this timing consistently from measurement to measurement (Methods). The timings of *Tn* and *Tz* were well-correlated with each other in both growth conditions (Fig. 2A,B) with a Pearson correlation *R* = 0.77 for moderately fast and *R* = 0.84 in slow-growth conditions. However, at least part of this correlation can be explained by the fact that FtsZ protofilament assembly is required to recruit FtsN. At the same time, there was a significant lag time from the onset of Z-ring formation to the onset of FtsN recruitment (Fig. 2C,D). The average lag time (all averages will be denoted by <>) was about a quarter of the cell cycle time for both media conditions (Glucose-cas: < (*Tn* – *Tz*)/*Td* >= 27 ± 17%; glycerol-TrE: 27 ± 15%) (SI Table S1). This long lag time shows that midcell formation of a persistent Z ring alone is not sufficient for the recruitment of FtsN.

**Figure 2.**
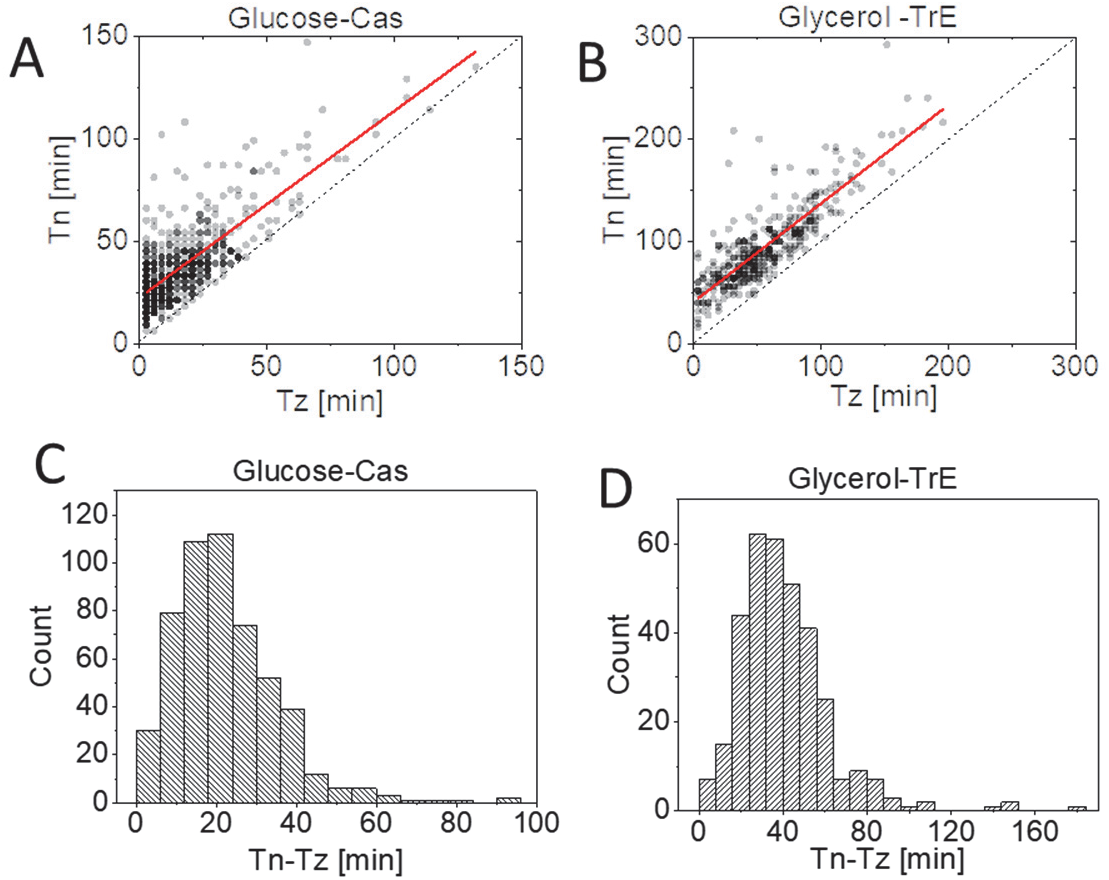
The time delay between the FtsZ-ring/ZapA and N-ring formation. **A-B** Timing of the formation of a persistent Z-ring, *Tz*, versus timing for midcell accumulation of FtsN, *Tn*, for cells grown in M9 minimal media supplemented with **(A)** glucose-cas (N=526) and **(B)** glycerol-TrE (N=339). The solid red lines are the linear fit to the data (*Tn* = (0.91 ± 0.03)*Tz* + (22 ± 0.8*min*), *R* = 0.77) for glucose-cas and *Tn* = 0.96 ± 0.03*Tz* + (41 ± 2.2*min*), *R* = 0.84) for glycerol-TrE cells. The dashed black line corresponds to *Tn* = *Tz*. **C-D** Distributions of time delays between the Z-ring and N-ring formation (*Tn* – *Tz*) for cells grown in **(C)** glucose-cas media (21 ± 13*min*; *mean* ± *SD*) and **(D)** in glycerol-TrE media (39 ± 22*min*; *mean* ± *SD*).

### ZapA fraction at midcell reaches a maximum at the start of the FtsN recruitment to the divisome

Why is the midcell recruitment of FtsN delayed by about a quarter of the cell cycle, and what event is needed for the recruitment of FtsN to occur? It has been argued that the onset of constriction in *B. subtilis* follows condensation of FtsZ protofilaments in the middle of the cell (40, 45). To test if a similar process also holds in *E. coli*, we determined the width of the Z-ring and the N-ring at the time of the onset of constriction (*Tn*). The width reflects the spatial spread of FtsZ protofilaments along the long axis of the cell, however, the number reported is larger than the actual spread because of the width of the point-spread function (PSF) of the microscope. In both growth conditions, the width of midcell ZapA-mCherry accumulations started to decrease before the onset of constriction and continued to decrease throughout the constriction period (Fig. 3A,B). The width of the N-ring also decreased throughout the constriction period. Thus, there was no sharp condensation in the longitudinal distribution of ZapA/FtsZ before or at the onset of constriction, which is not consistent with an abrupt FtsZ protofilament condensation needed to trigger the onset of constriction.

**Fig. 3.**
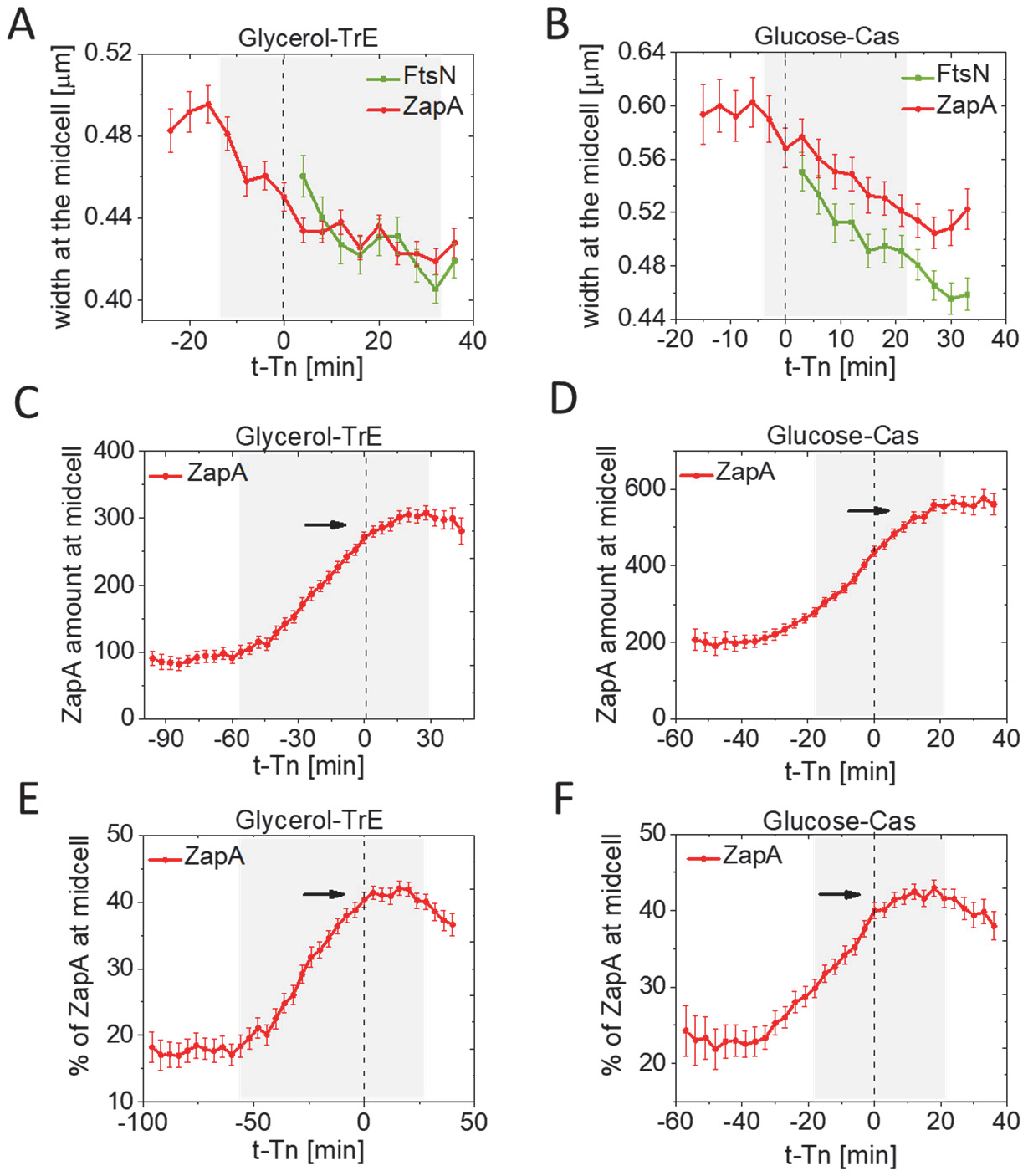
Kinetics of ZapA accumulation at the time of FtsN recruitment at midcell. **A-B** The spatial spread of ZapA-mCherry and Ypet-FtsN accumulations at midcell along the long axes of a cell as a function of time. Time zero corresponds to *Tn* (indicated by a dashed vertical line). **(A)** Slow-growing cells in glycerol-TrE media (N=339) and **(B)** moderately fast-growing cells in glucose-cas media (N=526). The shaded area marks the region where the number of cells analyzed is no less than 10% of its maximal value. All error bars represent 95% confidence intervals. **C-D** Fluorescent intensity of ZapA-mCherry at midcell (1.0 μm wide band) as a function of time from FtsN recruitment at midcell for **(C)** slow-growing and **(D)** moderately fast-growing cells. The fluorescent intensity reflects the number of ZapA present at the midcell. The arrow points to a plateau region. **E-F** The percentage (fraction) of ZapA at midcell (1.0 μm wide band) as a function of time from FtsN recruitment at midcell for **(E)** slow-growing and **(F)** moderately fast-growing cells. The arrow points to a plateau region.

It has also been proposed earlier that FtsZ needs to accumulate to some threshold number at the Z-ring to initiate cell division (46). To test this idea further, we determined how the number of ZapA molecules varied as a function of time at about time *Tn*. In slow-growth conditions, the number of ZapAs in the divisome increased linearly in time before *Tn* (Fig. 3C). At time *Tn*, the linear increase almost stopped, and the number of ZapA molecules stayed constant for about 30 min (Fig. 3C). This behavior is consistent with the idea of a threshold accumulation of FtsZ. However, at a moderately fast growth rate, the number of ZapA molecules increased continuously throughout *Tn*, and their number peaked only about 20 min (0.25Td) after the onset of constriction (Fig. 3D). This further increase could have arisen from fluorophore maturation if the concentration of ZapA-mCherry had varied within the cell cycle, but we found this variation small (<10%).

We subsequently investigated how the fraction of ZapA at the midcell varies as a function of time in the vicinity of *Tn*. We found that the population-average fraction also increased about linearly after formation of a persistent Z-ring (Fig. 3E,F). At time *Tn*, the fraction stopped increasing in both growth conditions and plateaued at its highest value (43% in both growth conditions). So, the threshold accumulation of a relative *fraction* of ZapA holds in both growth conditions. The same behavior was also confirmed by plotting *Tn* vs this fraction and observing that these two quantities showed a zero correlation (SI Fig. S3). It is unclear why the fraction of ZapA should reach a threshold value rather than the total number of ZapA molecules in the divisome at the onset of constriction. Furthermore, the coefficient of variation for both the fraction of ZapA and the total number of ZapAs in the midcell Z-ring was about 0.3 (SI Fig. S3), indicating that if there is a threshold, then it is rather poorly defined in the cell population. An alternative explanation for plateauing is that at time *Tn* some change in the divisome prevents a further increase in the FtsZ amount in the Z-ring. Rather than being a cause for the onset of constriction, the threshold accumulation may be a consequence of divisome maturation.

### Septal cell wall synthesis starts at the first distinct stage of the FtsN recruitment

Our single-cell data (cf. Fig. 1B, middle) is indicative that the recruitment of FtsN occurs in two stages during the cell cycle, although single-cell signals show large fluctuations. This finding would support the idea that FtsN is first weakly recruited to the divisome via FtsA and later more strongly via its SPOR domain once septal PG synthesis starts (16, 30, 33). To investigate this idea further, we averaged the single-cell data over the cell population to remove the large fluctuations in protein numbers inherent to single-cell data. The population-averaged midcell accumulation of FtsN showed two distinct stages when plotted as a function of cell age (Fig. 4A,B). The first rise in the FtsN midcell amount occurs at about the population-averaged time <*Tn*>, which is (0.48 ± 0.11)*Td* in glucose-cas and (0.65 ± 0.14)*Td* in glycerol-TrE measurements (*mean* ± *SD*). Expectedly, the sharp increase observed in the single-cell data (cf. Fig. 1B middle) was smeared out because of the variability in *Tn* timing in the cell population. There was also a second distinct increase in FtsN accumulation in the population-averaged data. It started at about 0.75*Td* for cells in the glucose-cas (Fig. 4A) and 0.9*Td* for cells in the glycerol-TrE medium (Fig. 4B). This increase was accompanied by a *decrease* in the relative and absolute amount of ZapA/FtsZ in the Z-ring. An increase in the FtsN midcell numbers, while there was a decrease in the ZapA/FtsZ numbers, indicates FtsN is recruited independently of ZapA/FtsZ during this stage. Such recruitment may be due to binding of the FtsN SPOR domain to denuded glycan strands and not involving ZapA/FtsZ.

**Fig.4.**
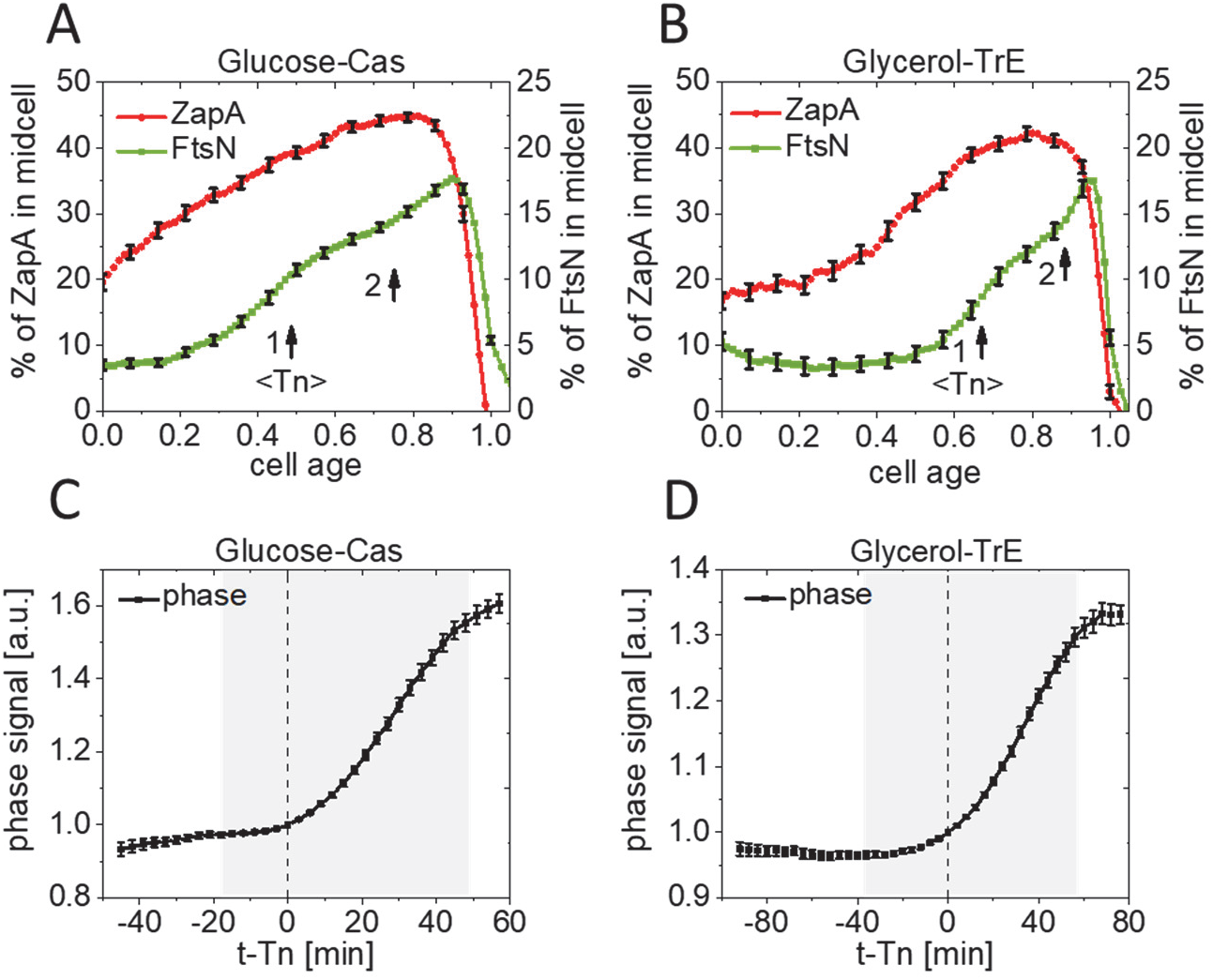
Septal cell wall synthesis starts at the first distinct stage of FtsN recruitment. **A-B** The percentage/fraction of ZapA-mCherry (left axis) and Ypet-FtsN (right axis) at midcell (1.0 μm wide band) as a function of cell age for **(A)** moderately fast-growing (glucose-cas media, N=526), and **(B)** slow-growing cells (glycerol media, N=339). Arrows show the approximate start of stages 1 and 2. The start time for stage 1 is <*Tn*> which is based on data in Fig. 2 and Table S1. The error bars represent 95% confidence intervals (for clarity, only every 5th point is shown). **B-C** Phase signal intensity at midcell (1.0 μm wide band) as a function of time from FtsN recruitment at midcell for **(C)** moderately fast-growing and **(D)** slow-growing cells. Time zero corresponds to *Tn* (indicated by a dashed vertical line). The shaded area marks the region where the number of cells analyzed is no less than 10% of its maximal value. All error bars correspond to 95% confidence intervals.

It is also worth mentioning that only a relatively small fraction of FtsN is recruited to the divisome even at its peak recruitment. At the peak during the 2nd stage, we found that the number of FtsN in the divisome was just 16% of the total number of FtsN molecules present in the cell in glycerol-TrE and 18% in glucose-cas medium (Fig. 4A,B; SI Fig. S4). In contrast, about 45% of ZapA is recruited to the divisome at its peak recruitment. It is unclear what the function of the remaining 83% of the FtsN molecules, which appear to be uniformly distributed throughout the cell membrane. Of the 17% of the FtsN present at midcell at its peak, 70% were recruited during the first stage, while the remaining 30% were recruited at the second stage. Interestingly, these fractions were the same in both growth conditions. Moreover, if the recruitment kinetics are plotted as a function of (absolute) time instead of cell age (relative time), then the duration of the first and the second stages were effectively the same (SI Fig. S5A,B) despite the twofold difference in doubling times in these two growth conditions (SI Table S1). The latter finding is suggestive that septal PG synthesis is not limited by substrate availability, which is likely to vary in different growth conditions, but by enzyme kinetics. Next, we investigated how the two stages of FtsN recruitment are related to septal cell wall synthesis. We used two independent methods to assess the synthesis rate of the septal cell wall. In the first method, we followed the phase signal at midcell. The change in the latter is approximately proportional to the change in the amount of the dry biomass at the midcell (see Methods). As the septum starts to constrict, the dry biomass in the cell center decreases, leading to an increase of the phase signal in the negative phase contrast imaging that we used (cf. Fig. 1B, bottom). The increase in the phase signal approximately coincided with the onset of the first stage of the FtsN recruitment (within about ±3*min* glucose-Cas and ±8*min* in glycerol medium) when we aligned phase signals of single cells relative to their *Tn* (Fig. 4C, D). As a second method, we determined the length growth rate of the cells. Here the length growth rate is defined as (1/*L*)(*ΔL*/*Δt*) where *ΔL* is the change of the cell length from a pole to a pole, *L*, during small time interval *Δt*. This growth rate varies in a cell-cycle-dependent manner rather than being a constant as shown recently (47). The length growth rate also started to increase when the first FtsN accumulation occurred in the glycerol-TrE medium (SI Fig. 5E,G). In glucose-cas medium, the length growth rate may be delayed but not more than by about 0.1Td (SI Fig. 5F,H). These two independent measurements thus show that septal PG synthesis starts already at the first stage of FtsN recruitment to the divisome.

**Fig.5.**
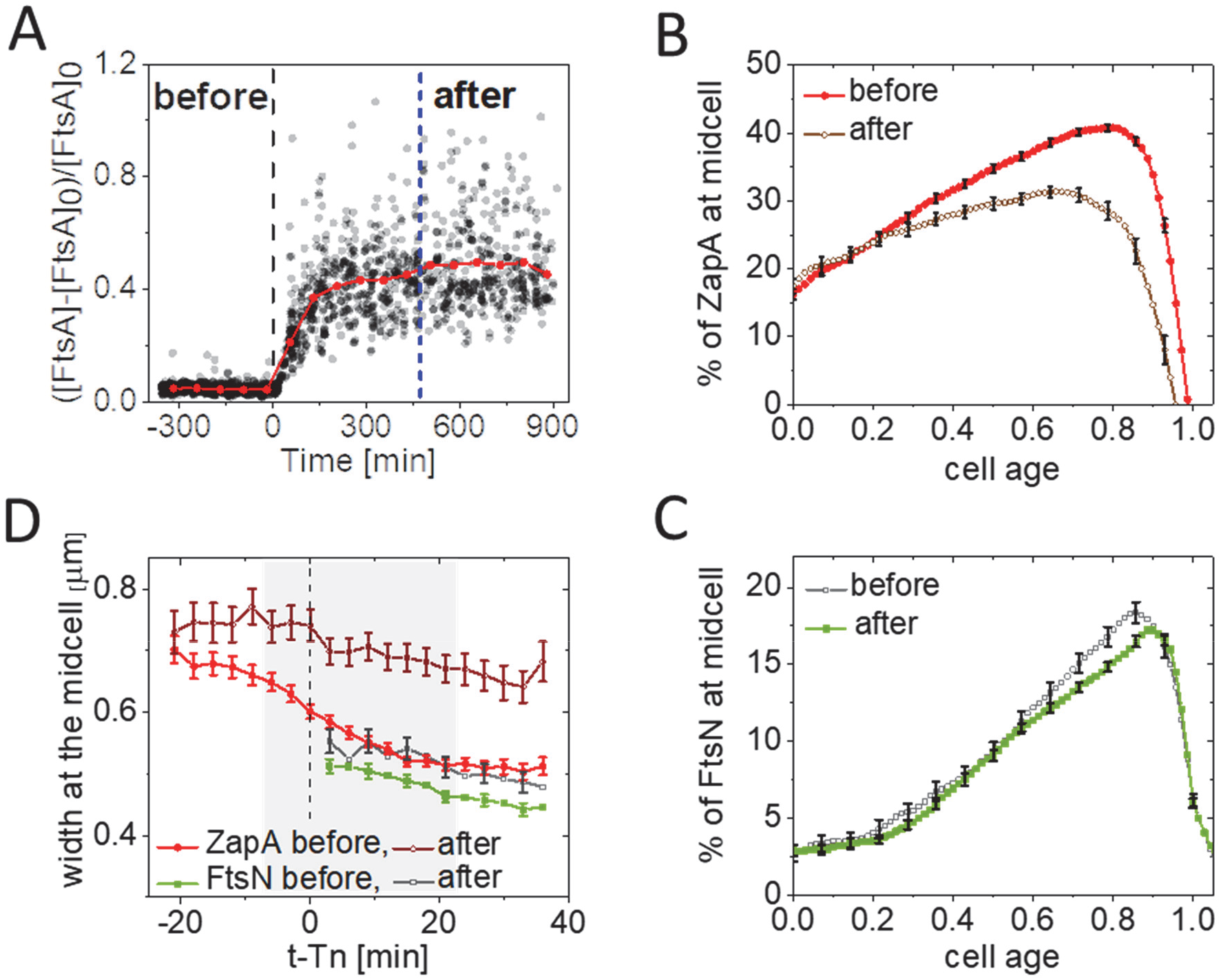
Changes in FtsN and ZapA midcell accumulations upon FtsA overexpression. **(A)** A relative increase in the concentration of FtsA, [FtsA], at cell birth before and after induction with 100μM of IPTG as inferred from monitoring GFP reporter expressed from pDSW210 (strain JM149). The shown increase is relative to WT FtsA concentration, [FtsA]0, which is determined by Western blotting. Time zero (black dashed vertical line) corresponds to the start of the induction. N=1470. **(B)** The percentage (fraction) of ZapA-mCherry at midcell before and after overexpression of FtsA as a function of cell cycle time. For cells termed “after,” ZapA amounts were analyzed when the cell growth reached a new steady-state (indicated by a blue vertical dashed line in panel A). Cells were grown in M9 glucose-cas media at 28°C. The error bars represent 95% confidence intervals (for clarity, only every 5th point is shown). N=660 (before); N=223 (after). **(C)** The percentage of Ypet-FtsN at midcell before and after overexpression of FtsA as a function of cell cycle time. Conditions as above for ZapA-mCherry in (B). **(D)** The spatial spread of ZapA-mCherry and Ypet-FtsN accumulations at midcell along the long axes of cell as a function of time before and after overexpression of FtsA. Midcell traces of ZapA-mCherry and Ypet-FtsN were aligned at the time of FtsN recruitment at midcell, *t – Tn* = 0, marked by a dashed line. The shaded area marks the region where the number of cells analyzed does not vary more than 10%. The error bars represent 95% confidence intervals (for clarity, only every 2nd point is shown for FtsN).

### Upregulation of FtsA affects recruitment of ZapA/FtsZ and condensation of the Z-ring but not the N-ring

To further understand what role FtsA plays in the recruitment of FtsN and early divisome proteins, we introduced into our strain an extra copy of FtsA on plasmid pDSW210 (pSEB306+) under an isopropyl-β-D-thiogalactoside (IPTG) inducible P_trc_ promoter (48, 49). This plasmid allows upregulation of the concentration of FtsA ([FtsA]) in the cell by about 50% (SI Fig. S6) under the experimental conditions used in this study. However, in slow growth conditions in glycerol medium, FtsA expression from this plasmid is too leaky (SI Fig. S6B), so we focus on the results from the moderately fast growth condition. In addition to the strain mentioned above, we constructed a reference strain expressing GFP instead of FtsA from pDSW210 in non-labeled WT background BW27783 cells. We imaged this reference strain (JM149) in the same microfluidic chip simultaneously with the above-described strain of interest (JM150). The total fluorescence intensity of GFP in the reference strain allowed us to estimate the number of FtsA molecules synthesized from plasmid pSEB306+ before and after IPTG induction revealing that leakage from the plasmid accounted for less than 5% of the native protein level in uninduced conditions (Fig. 5A). The induced condition resulted in a 50% increase in the level of FtsA, which led to an increase in the cell length by about 10% (SI Table S1). This behavior is different from the high-level expression of FtsA that inhibits cell septation and causes the formation of filamentous cells (50–52).

The main effect observed after overexpression of FtsA was a significantly reduced amount of ZapA-mCherry at the divisome (Fig. 5B). However, in the first quarter of the cell cycle, the amount of ZapA-mCherry at midcell was not affected. During the first quarter, ZapA-mCherry is mostly part of transient FtsZ assemblies and other complexes that lack FtsN. In later parts of the cell cycle, the midcell ZapA fraction decreases from about 45% to 30% at its peak. At the same time, strikingly, the fraction of FtsN at midcell remained unaffected by FtsA upregulation throughout the cell cycle (Fig. 5C). Thus, the FtsA upregulation by 50% does not affect recruitment of FtsN to midcell, but it has a significant effect on ZapA/FtsZ recruitment. We also investigated how the increased FtsA affected the condensation of the Z-ring. Based on *in vitro* measurements, FtsA has been suggested to act as an anti-bundling agent through forming minirings (53, 54). We, therefore, compared the changes in the width of both the Z-and N-rings before and after the onset of constriction (*Tn*). Before FtsA induction, both the Z-and N-ring widths behaved similarly to the WT strain without the plasmid (Fig. 5D). However, in FtsA upregulated conditions, the distribution of FtsZ protofilaments along the long axes of the cell remained wide and did not condense. In contrast, the N-ring appeared similar to WT cells (Fig. 5D). This finding indicates that FtsN and ZapA/FtsZ are present in spatially separated complexes. These conclusions agree with previous observations that proteins in the Z-ring were separated from proteins in the PG synthesizing machinery along the radial direction (19, 55, 56). Here, our data show that separation can also occur along the cell’s long axis, at least in FtsA upregulated conditions. Furthermore, the finding that the ZapA-mCherry (FtsZ)-ring stays broad and does not condense suggests that FtsA upregulation antagonizes FtsZ protofilament bundling. Strikingly, however, the recruitment of FtsN and the condensation of the N-ring were not affected. The latter finding suggests that the majority of FtsN is recruited to the divisome independent of ZapA/FtsZ.

### FtsA* leads to earlier recruitment of FtsN but not to earlier constriction

It has been proposed that FtsA self-interaction prevents recruitment of downstream proteins, including FtsN, to the divisome (31, 32, 57). The FtsA mutant R286W (FtsA*) has a reduced ability to self-interact/polymerize but still interacts normally with FtsZ (57–59). To further understand how disruption of the FtsA self-interaction affects FtsN recruitment kinetics, we upregulated FtsA* in the WT FtsA background. To that end, we replaced *ftsA* by *ftsA*\* in plasmid pDSW210 (pSEB306+*) and repeated experiments described in the previous section. We found that upregulation of FtsA* by about 50% with respect to the native FtsA levels did not affect cell doubling times but led to about a 16% shortening of cell length (SI Table S1). Overexpression of FtsA* decreased the midcell fraction of ZapA-mCherry slightly (Fig. 6A). The decrease of about 3% was uniform throughout the cell cycle in contrast to overexpression of FtsA, in which case the ZapA fraction in the beginning of the cell cycle was not affected but in the late stages decreased by about 15%. Furthermore, we found that the width of the midcell Z-ring was not affected (SI Fig. S7A). At the same time, significantly more Ypet-FtsN was recruited at midcell following the overexpression of FtsA* and it arrived at the same time as ZapA. Importantly, the fraction of FtsN at midcell was almost uniformly increased throughout the cell cycle (Fig. 6B). It was noticeable that the midcell fraction of FtsN in the FtsA* upregulated conditions increased throughout the cell cycle in the same way as the midcell fraction of ZapA, although the latter was about twice higher (Fig. 6C,D). Unlike FtsA, FtsA* is thus capable of recruiting FtsN to the divisome as soon as the Z-ring is assembled and in proportion to the fraction of ZapA/FtsZ in the Z-ring.

**Fig.6.**
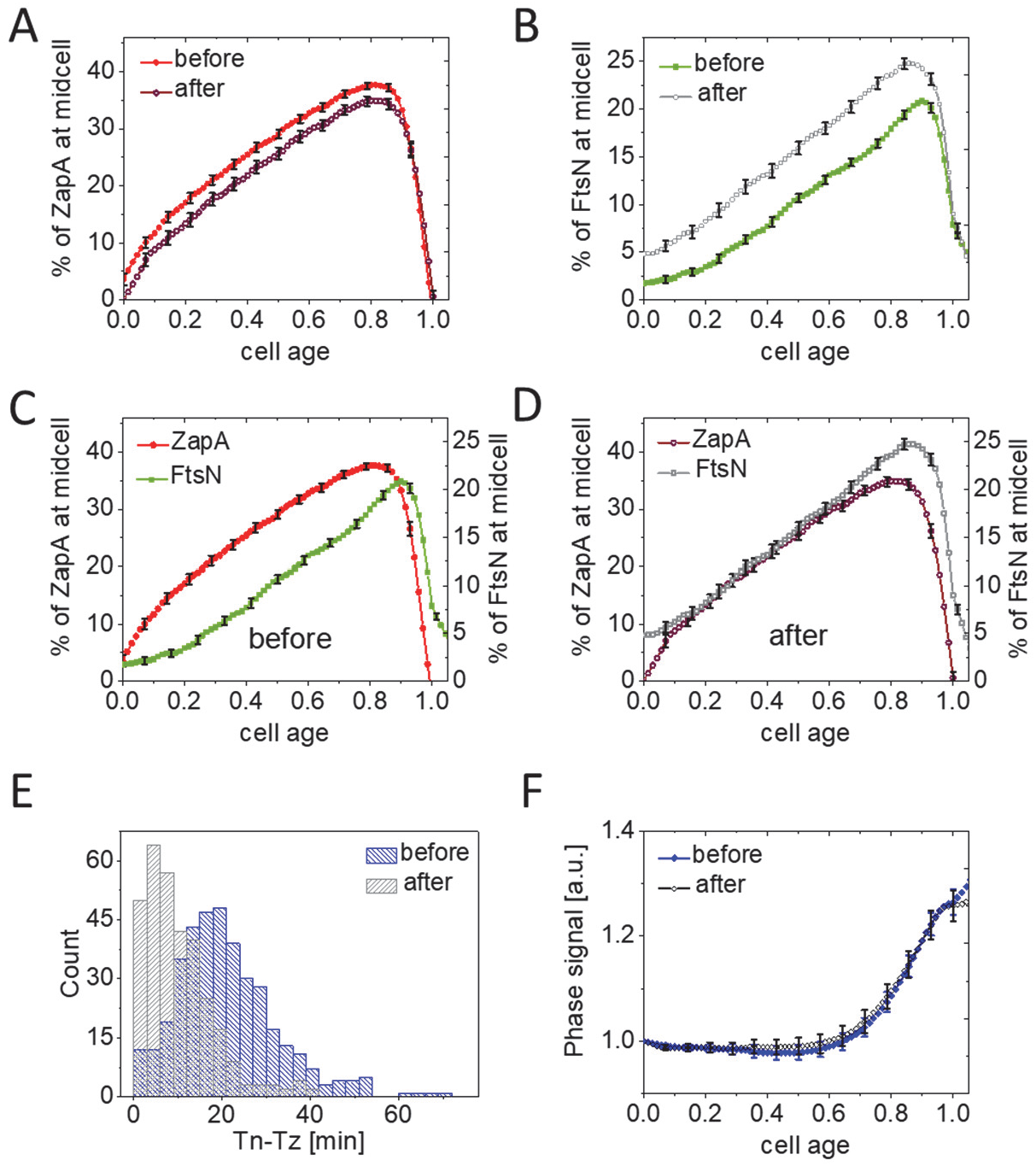
Changes in FtsN and ZapA midcell accumulations upon FtsA* (FtsA^R286W^) overexpression. **(A)** The percentage (fraction) of ZapA-mCherry at midcell before and after overexpression of FtsA* (R286W) as a function of cell cycle time. FtsA* was expressed as an extra copy from a plasmid pSEB306+*by the addition of 100μM IPTG in M9 glucose-cas media at 28°C. For cells termed “after,” ZapA amounts were analyzed when the cell growth reached to a new steady-state. All error bars represent 95% confidence intervals (for clarity, only every 5th point is shown). N=381 (before); N=321 (after) **(B)** The percentage of Ypet-FtsN at midcell before and after overexpression of FtsA* as a function of cell cycle time. Conditions as above for ZapA-mCherry in (A). **(C)-(D)** The percentage of ZapA-mCherry and Ypet-FtsN at midcell before and after upregulation of FtsA*, respectively. **(E)** The distributions of time delay between the Z-ring and N-ring formation (*Tn – Tz*) in cells before (20 ± 12; *mean* ± *SD*) and after (9 ± 8; *mean ± SD*) upregulation of FtsA*. **(F)** Phase signal intensity at midcell (1.0 μm wide band) as a function of cell age before and after upregulation of FtsA*.

In addition to higher midcell abundance, FtsN recruitment also shifted earlier (from 34 ± 18 *min* to 26 ± 17 *min*; t-test p=4.1×10^−10^), and the delay between recruitment of ZapA and FtsN to midcell shortened from 20 ± 12 *min* without FtsA* to 9 ± 8 *min* in its presence (Fig. 6E). The distribution of these delay times changed from one with a distinct lag time to an exponential, indicating that the presence of FtsA* eliminates the lag. However, the earlier and more abundant recruitment of FtsN to the divisome did not lead to the earlier onset of constriction in these conditions (Fig. 6F, SI S7B). This result indicated that the recruitment of FtsN to the divisome is not a rate-limiting process for the onset of constriction under these conditions but reveals that FtsA* accelerates FtsN recruitment.

## Discussion

FtsN has been implicated as a trigger for the onset of constriction (16, 18, 20). Here we studied its recruitment kinetics to the divisome using an early cell division protein ZapA as a reference after verifying that the latter is a good proxy for FtsZ. Our data show that, unlike the gradual increase of ZapA/FtsZ at midcell over about a quarter of the cell cycle, the recruitment of FtsN occurs abruptly over much shorter time scales and, on average, about a quarter of the cell cycle later than when the persistent Z-ring forms. The question then arises as to what is responsible for the delay in FtsN recruitment following formation of the Z-ring? A long delay between the formation of a persistent Z-ring (*Tz*) at midcell and an abrupt accumulation of FtsN (*Tn*) suggests that there is some cell cycle checkpoint between the formation of a persistent Z-ring and the onset of constriction.

### Is the onset of constriction triggered by the change in FtsZ protofilament assembly at midcell?

What could be responsible for the delay between *Tz* and *Tn*? The putative mechanisms proposed for a checkpoint for *Tn* so far include the change in the polymerization state of FtsZ (46), its higher-order assembly (40, 45) or that of FtsA (31, 32) or simply recruitment of FtsN itself (17). To further investigate the idea that FtsZ dynamics drive the checkpoint, we followed the amount of ZapA/FtsZ and the effective width of the septal ring before and after the onset of constriction. Our data show that condensation of FtsZ protofilament assemblies started before the onset of constriction and continued well past it. Unlike *B. subtilis* cells (39, 40), we did not observe a change in FtsZ protofilament condensation at the onset of constriction. It is possible that the mechanism triggering the onset of constriction is different in these two organisms, although many of the essential divisome components are homologous to each other.

We also tested the hypothesis that the numbers of FtsZ molecules need to reach some threshold value at midcell to initiate the formation of a constriction (46). Our data show that ZapA/FtsZ reaches a broad maximum (plateau region) in slow growth conditions at the onset of constriction. However, in faster growth conditions, this maximum is reached later. On the other hand, the fraction of ZapA/FtsZ at midcell (about 45%) plateaued at the onset of constriction in both growth conditions. It is unclear why a fraction of FtsZ at midcell could have any significance in initiating downstream biochemical processes. A more likely explanation is that the constant fraction of ZapA/FtsZ at midcell indicates that divisome has matured to a functional form for PG synthesis. This mature divisome appears to be in a quasi-equilibrium characterized by an equilibrium constant. As the cell syntheses more ZapA/FtsZ, it is incorporated into a divisome in proportion to this equilibrium constant. The constant fraction thus may be the consequence of divisome maturation by some unrelated process rather than the cause, but further work is warranted to support this claim.

### Recruitment of FtsN is not rate-limiting for the onset of constriction

Our data shows that there is sufficient FtsN available at all cell cycle times, but only about 17% is present in the septal ring at its peak, which occurs in the late stages of cell division (0.1-0.15Td before division). The late recruitment of FtsN to the divisome is thus not because FtsN is not available. This assessment is also consistent with measurements where we expressed FtsA* in addition to the native copy of FtsA. In these conditions, FtsN was recruited concurrently with the formation of the Z ring. The early recruitment of FtsN in this strain can be explained by an increased number of FtsA* molecules with a free 1C domain to interact with FtsN since FtsA* is defective in self-interaction, which would mask the 1C domain (57–59). Though FtsN is available early in the cell cycle in WT cells, its recruitment does not occur until a later stage of the cell cycle. This indicates that some other process holds back the maturation of the divisome and that recruitment of FtsN to the divisome is not a rate-limiting process for the onset of constriction. It is possible that instead of FtsN, the recruitment of one or more of FtsN’s interaction partners is rate-limiting. The potential interaction partners that could be rate-limiting for division are FtsQ (in complex with FtsBL) or FtsWI proteins (28, 60). It could also be that the rate-limiting step is a change in FtsA polymerization/self-interaction state or the activation of FtsW/I (57, 61). Although data show that the recruitment of FtsN is not rate-limiting for the onset of constriction, it does not contradict the notion that recruitment of FtsN is the essential last step that triggers activation of the divisome. Instead, our findings show that the recruitment of the FtsN is not instrumental in controlling the timing of the division process, as it should occur rapidly once the rate-limiting step is surmounted.

### Decoupling of FtsN midcell accumulations from FtsZ protofilaments

Our data (Fig. 5) show that after FtsA overexpression, the fraction of ZapA/FtsZ in the Z-ring decreased, and the rings did not condense at the onset of constriction as it did in WT cells. This mild effect of increased FtsA on Z ring morphology is consistent with the disassembly of Z rings that occurs when FtsA is increased 10-fold (50). What is surprising is that the N rings assembled normally. This finding can be explained if FtsN in the septal ring is not physically linked to FtsZ protofilaments. Evidence for decoupling FtsN and FtsZ protofilaments has also been documented in other measurements where their localization in the radial direction was examined (55) and from recent single-molecule studies (62).

Beyond ZapA/FtsZ and FtsN being part of different complexes, the data on Fig. 5 furthermore could imply that FtsA has a minor role in directly recruiting FtsN to the septal ring. Otherwise, upregulation of FtsA would have either up or downregulated the FtsN fraction in the midcell as it did for the ZapA/FtsZ fraction and as it occurred when FtsA* was upregulated. However, this interpretation might be too simplistic because it assumes that FtsA forms similar polymeric or higher-order structures when associated with FtsZ protofilaments and in separate divisome complexes together with FtsN. If this is not the case, then the two different higher-order structures could respond differently to increasing FtsA concentration. When in complex with FtsZ protofilaments, FtsA could form 12 member minirings as observed *in vitro* measurements (53, 54, 63), which appear to prevent the condensation of FtsZ protofilaments. As such, FtsA acts as an anti-bundling agent for FtsZ protofilaments, which is consistent with findings in Fig. 5 that upregulation of FtsA leads to more diffuse Z-rings and a lower fraction of ZapA/FtsZ in these rings. The concentration of the minirings should have a sensitive dependence on the total FtsA monomer concentration with a high Hill coefficient (^~^12). As a result, small variations in FtsA concentration could significantly affect FtsZ protofilament assemblies observed in our experiments.

A different scenario might materialize in divisome complexes containing FtsN. FtsN and potentially other divisome proteins can be expected to bind to FtsA via its 1C interface (29, 57, 64, 65). In support of his assumption, when FtsA* is present, FtsN is rapidly recruited to the Z ring overcoming the normal delay observed when only WT FtsA is present. The 1C domain is also involved in forming a monomer-monomer interface in FtsA polymers (66). Because of the dual role of this interface, FtsA cannot form minirings when it is already a part of the divisome complex as the latter prevents the closure of the ring. Thus, active divisome complexes should only include linear FtsA polymers. The length of these polymers could be rather short because linear polymers should be less stable than the ring ones. Furthermore, the length distribution of linear polymers can be expected to have a weaker dependence on the total FtsA monomer concentration. A weaker dependence would be consistent with the experimental observation that FtsN concentration at midcell does not change upon upregulation of FtsA (Fig. 5C). So far, there is no experimental data on the polymerization state of FtsA in either of the complexes. It remains even a possibility that divisome complexes with FtsN do not contain any FtsA. Furthermore, the FtsA minirings have not been observed *in vivo*, and it remains unclear how the minirings could keep up with treadmilling FtsZ protofilaments. Further measurements are warranted to understand what is the polymerization state of FtsA in different complexes *in vivo*.

### Two different stages in septal peptidoglycan synthesis

Our data reveal two distinct stages in FtsN recruitment to midcell, both involving septal PG synthesis. We observed that the first FtsN recruitment/accumulation stage coincides with the onset of the constriction/ septal PG synthesis within about ± 4 min in glucose-cas and ± 8 min in glycerol medium (Fig. 1A; 4C-D). This may imply that the arrival of FtsN results in rapid activation of septal PG synthesis, although based on the time-resolution of our measurements, it is also possible that initial septal PG synthesis starts first and FtsN is recruited thereafter. In the former scenario, FtsN could be recruited first via its binding to FtsA (29, 30, 32). However, in this case, there is no extended period when FtsN lingers in the divisome just due to its sole binding to FtsA because once septal PG synthesis starts, FtsN can bind to denuded peptidoglycan strands also via its SPOR domain. This is presumably stronger binding than the binding of FtsN to FtsA, but the latter is not neglible because cells without the FtsN cytoplasmic domain or the FtsN D5N mutants show a division defect and elongated phenotype (31, 32). The second distinctive stage of FtsN accumulation occurs late in the cell cycle (15 min before division in both growth conditions corresponding to 0.81Td and 0.90Td for glucose-cas and glycerol conditions, respectively). There seems to be some additional mechanism involved in this stage, in addition to the SPOR-domain related self-enhanced recruitment of FtsN. Interestingly, the start of the second stage of FtsN recruitment occurs approximately when the amount of ZapA/FtsZ in the septal ring starts to decrease. While this could be a mere coincidence, it appears that FtsZ protofilament-involved complexes may sequester away some of the FtsN binding partners such as FtsQLB from the divisome complexes. Once FtsZ protofilaments at midcell become unstable, perhaps because of high membrane curvature at the site of constriction, fewer FtsQLB complexes are associated with FtsZ protofilaments, and more can be directly recruited by the divisome complexes. As the elevated level of FtsN at the septal ring coincides with the increase in septal cell wall synthesis (SI Fig. S5, C-F), the rate of the septal cell wall synthesis increases in this second stage (SI Fig. S5 C-D). At the very end of the division process, this synthesis proceeds without any FtsZ protofilaments present at the division site. It is likely that FtsN and, to a lesser degree DedA, which also features a SPOR domain (67), completely take over the role of positioning the divisome apparatus from FtsZ protofilaments.

The two phases of FtsN accumulation may also correspond to the septation stages recently observed in TEM images of dividing *E. coli* (68). The images showed a V-shaped invagination in the early stages of septation, referred to as the constriction phase. This invagination developed an inward protrusion similar to the septum of *B. subtilis* in the second stage, referred to as the septation phase. Although the authors did not determine the exact timing of these phases in the cell cycle, their approximate durations appear similar to stages one and two observed here for FtsN recruitment.

In conclusion, our data show an abrupt accumulation of FtsN to the divisome, which occurs about 30% of cell cycle time after a persistent Z-ring forms. This accumulation coincides with the onset of septal cell wall synthesis within the uncertainties of our measurements. We did not observe a change in FtsZ protofilament condensation at the start of constriction but did observe that the fraction of FtsZ in the ring reached a plateau. Furthermore, our data support the idea (55, 62) that FtsN is not part of the complexes that include FtsZ protofilaments but resides in separate divisome complexes involved in septal PG synthesis. Finally, our data show that the recruitment of FtsN to midcell is not rate-limiting for the onset of constriction, even though this recruitment step is the last essential step for the onset of constriction.

## Experimental procedures

### Media, bacterial strains, and plasmids

Cells were grown with M9 minimal media (Teknova Inc., CA) supplemented with 0.5% glucose (Millipore Sigma, MO) and 0.2% casamino acids (cas, ACROS Organics) or 0.3% glycerol (ThermoFisher Scientific) and trace metals mixture (TrE, Teknova Inc., CA) at 28°C. Unless indicated otherwise, antibiotic concentrations were as follows: Ampicillin (Amp), 100 μg/mL; Chloramphenicol (Cm), 25 μg/mL; Kanamycin (Kan), 25 μg/mL). For induction, 100μM of IPTG was used. All *E. coli* strains used in the reported experiments are derivatives of K12 BW27783 obtained from the Yale Coli Genetic Stock Center (CGSC#: 12119). The strain with fluorescent fusions of ZapA-mCherry and Ypet-FtsN (JM144) was made by P1 transduction (69). All strains and plasmids used in this study are listed in SI Tables S2A and S2B, respectively.

Further details for cell preparation and culture in microfluidic devices, fluorescence microscopy, image analysis and Western blot analysis can be found in SI Text.

## Acknowledgments

The authors thank Ariel Amir and Sriram Tiruvadi Krishnan for useful discussions, Da Yang and Scott Retterer for help in microfluidic chip making, and Thomas Bernhardt and Jie Xiao for bacterial strains. A part of this research was conducted at the Center for Nanophase Materials Sciences, which is sponsored at Oak Ridge National Laboratory by the Scientific User Facilities Division, Office of Basic Energy Sciences, U.S. Department of Energy. This work was supported by the US-Israel BSF research grant 2017004 (JM), by the National Institutes of Health awards GM127413 (JM) and GM29764 (JL).

